# FIN-Seq: Transcriptional profiling of specific cell types in frozen archived tissue from the human central nervous system

**DOI:** 10.1101/602847

**Authors:** Ryoji Amamoto, Emanuela Zuccaro, Nathan C. Curry, Sonia Khurana, Hsu-Hsin Chen, Constance L. Cepko, Paola Arlotta

## Abstract

Thousands of frozen, archived tissues from postmortem human central nervous system (CNS) are currently available in brain banks. As single cell and single nucleus technologies are beginning to elucidate the cellular diversity present within the human CNS, it is becoming clear that transcriptional analysis of the human CNS requires cell type specificity. Single cell and single nucleus RNA profiling provide one avenue to decipher this heterogeneity. An alternative, complementary approach is to profile isolated, pre-defined cell types and use methods that can be applied to many archived human tissue samples. Here, we developed FIN-Seq (<u>F</u>rozen <u>I</u>mmunolabeled <u>N</u>uclei <u>Seq</u>uencing), a method that accomplishes these goals. FIN-Seq uses immunohistochemical isolation of nuclei of specific cell types from frozen human tissue, followed by RNA-Sequencing. We applied this method to frozen postmortem samples of human cerebral cortex and retina and were able to identify transcripts, including low abundance transcripts, in specific cell types.

## INTRODUCTION

The human central nervous system (CNS) comprises an extremely diverse set of cell types. While this heterogeneous cellular composition has been appreciated since the work of early anatomists, it was not until recently, with the advent of single cell and single nucleus RNA sequencing, that different cell types of the adult human cerebral cortex and retina have begun to be defined at the molecular level (Cherry et al., 2018; Darmanis et al., 2015; Hodge et al., 2018; Lake et al., 2016; Lake et al., 2018; Liang et al., 2019; Lukowski et al., 2018; Peng et al., 2019; Phillips et al., 2018). These studies have identified at least 16 neuronal subtypes in the adult human cerebral cortex and 18 major cell types in the adult human retina. While these pioneering studies have started to highlight the heterogeneity of the adult human CNS, more fine-grained distinctions among cell types are likely present, and will become more apparent with increased numbers of cells profiled. Given such heterogeneity, gaining mechanistic insight into human CNS development, function, and disease will require transcriptional profiling at both single cell and cell type-specific resolution.

Transcriptional profiling of heterogeneous populations is feasible with either single cell RNA sequencing (Macosko et al., 2015; Shekhar et al., 2016; Tasic et al., 2016; Zeisel et al., 2015) or bulk RNA sequencing of purified user-defined cell types labeled either genetically or with dyes and antibodies (Arlotta et al., 2005; Heiman et al., 2008; Lobo et al., 2006; Molyneaux et al., 2015; Siegert et al., 2012; Telley et al., 2016). Single cell RNA sequencing has become essential for cataloguing molecularly distinct cell types in heterogeneous tissues such as the CNS. However, sampling the whole tissue for rare cell types, such as cone photoreceptors, is expensive as large numbers of single cells need to be profiled. Alternatively, bulk RNA sequencing of user-defined cell types allows for the acquisition of transcriptomes of rarer cell types; thus, avoiding sequencing of a large number of more abundant cell types. Acquisition of more transcriptomes via single cell RNA sequencing is accelerating the discovery of potential new markers that could be used to isolate specific, rare cell populations from cellularly diverse tissues. We aimed to develop a method that enables bulk RNA sequencing of specific cell types and extends to archived frozen tissue. Thousands of frozen human postmortem brain tissue samples, including those with disease, are readily available through brain banks, and they represent a crucial resource that is immediately available and largely untapped. The abundance of archived CNS tissue samples is crucial for profiling transcriptional changes in rare diseases, and it is also likely that the number of biological replicates needed in human studies is high because of the natural genetic variation present among individuals.

While whole-cell approaches are incompatible with flash-frozen CNS tissue, the nuclei from frozen tissue stay intact and can be profiled. In addition, nuclear RNA has been successfully used as proxy for the cellular transcriptome (Barthelson et al., 2007; Grindberg et al., 2013; Habib et al., 2017; Krishnaswami et al., 2016; Lake et al., 2016; Lake et al., 2017). Single nucleus RNA sequencing has indeed been used for unbiased profile of neuronal subtypes from frozen, archived human cerebral cortex tissue (Lake et al., 2016). However, a complementary technology to isolate and bulk sequence nuclear RNA of user-defined cell types from frozen human CNS tissue is lacking.

Here, we developed FIN-Seq (<u>F</u>rozen <u>I</u>mmunolabeled <u>N</u>uclei <u>Seq</u>uencing), a technology that combines nuclear isolation, fixation, immunolabeling, FACS, and RNA sequencing to obtain the gene expression profile of specific neuronal subtypes from frozen, archived human CNS tissue. While some antibodies such as those against NeuN and SOX6 are known to work with fresh tissue (Kozlenkov et al., 2018), a method to apply a wider range of antibodies against cell type specific markers is not available. With FIN-Seq, we isolated and profiled specific excitatory and inhibitory neuronal subtypes from frozen human cerebral cortex tissue and cone photoreceptors from the frozen human retina. Successful isolation of cone photoreceptors, which constituted roughly 2% of the whole retina, signified that rare populations could be reliably profiled from a frozen tissue sample. Interestingly, we also found that the nuclear transcripts captured with FIN-Seq represented more of the whole-cell transcripts compared to single nucleus sequencing. This is a novel, cost-effective technology that could enable deep transcriptional analysis of user-defined cell types from widely-available frozen human CNS samples.

## RESULTS

### FACS isolation of immunolabeled nuclei from frozen mouse brain samples

To test whether sequencing of specific nuclear RNA from frozen tissue is feasible, we first tested it using specific nuclear populations isolated from the frozen mouse neocortex. To this end, we modified previously-published protocols that use intracellular antibody staining to isolate specific cell types (Hrvatin et al., 2014; Molyneaux et al., 2015; Pan et al., 2011; Pechhold et al., 2009; Yamada et al., 2010). Intact cells cannot be dissociated from frozen tissue, so we developed a protocol to isolate fixed antibody-labeled nuclei and extract nuclear RNA (Figure 1a). Nuclei isolation eliminates the need for enzymatic dissociation, which induces aberrant activation of immediate early genes (Lacar et al., 2016). From the flash-frozen neocortex of P30 mice, we sought to isolate two populations of projection neurons, Corticofugal Projection Neurons (CFuPN) and Callosal Projection Neurons (CPN). In the adult mouse brain, BCL11B (also known as CTIP2) is largely expressed in CFuPNs in layer 5b and 6 and in sparse populations of interneurons. SATB2 is expressed by CPNs in all layers (Molyneaux et al., 2007) (Figure 1b). BCL11B and SATB2 expression are largely mutually exclusive, with a small population of layer 5 neurons expressing both markers (Harb et al., 2016; Molyneaux et al., 2015) (Figure 1b, **Layer 5, inset)**. Upon isolation by homogenization, nuclei were fixed, immunolabeled with antibodies against BCL11B and SATB2, and separated into two populations by FACS: SATB2^LO^BCL11B^HI^ (BCL11B^+^) and SATB2^HI^BCL11B^LO^ (SATB2^+^) (n=2 for each population) (Figure 1c-d). On average, we collected 55,215 BCL11B^+^ nuclei and 102,016 SATB2^+^ nuclei per biological replicate. These results indicate that this protocol could isolate intact nuclei that are immunolabeled with user-defined intranuclear antibodies.

**Figure 1:**
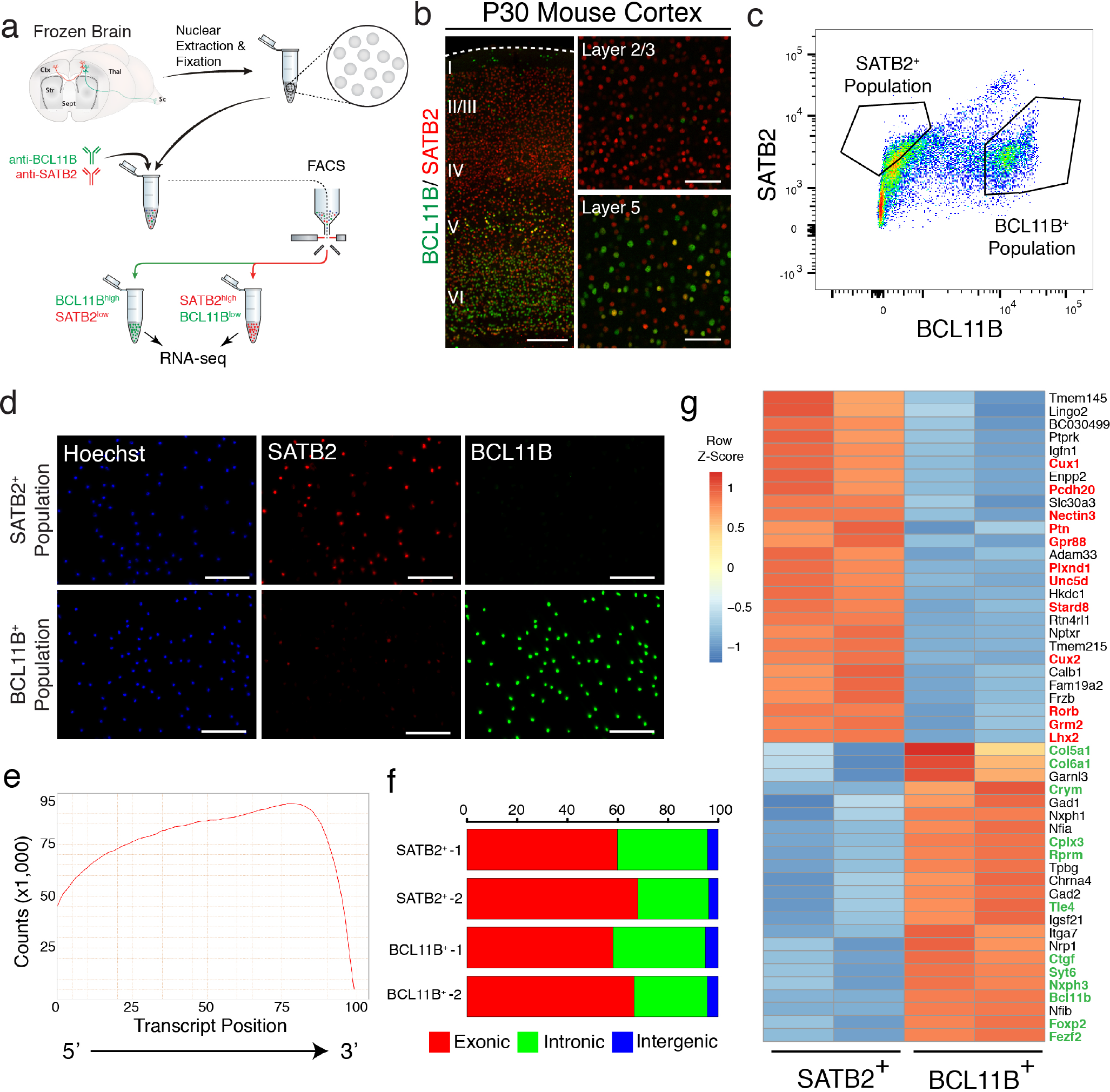
Isolation and transcriptome sequencing of two neuronal subtypes from the frozen mouse neocortex. (**A**) Schematic of FIN-Seq for frozen adult mouse brain. Nuclei were extracted from the frozen mouse neocortex by Dounce homogenization. The nuclei were fixed and immunolabeled with anti-BCL11B and anti-SATB2 antibodies. Two nuclear populations were isolated by FACS based on expression level of these two proteins. The nuclei were reverse crosslinked by protease digestion, and the RNA was extracted. Sequencing libraries were generated and subsequently sequenced to obtain cell type specific transcriptomes. (**B**) Representative immunohistochemistry images using BCL11B and SATB2 antibodies in the P30 mouse neocortex showed SATB2 expression in the upper layers and BCL11B expression in the deep layers (Left image). In layer 5, there were sparse cells that express both SATB2 and BCL11B (Bottom right image). (**C**) FACS plot of nuclei labeled with SATB2 and BCL11B antibodies showed a cluster of nuclei immunolabeled with BCL11B and a cluster of nuclei labeled with SATB2. (**D**) Isolated nuclei were counterstained by the Hoechst dye and either SATB2 or BCL11B in the SATB2^+^ population (top panels) or BCL11B^+^ population (bottom panels). (**E**) Representative quantification of read counts mapped by transcript position (5’ to 3’) for every gene. (**F**) Representative quantification of percentage of read counts mapped to exonic, intronic, or intergenic regions of the genome. (**G**) Heatmap of unbiased top 50 differentially expressed genes between SATB2^+^ and BCL11B^+^ populations. Known markers of callosal projection neurons (in red) were enriched in the SATB2^+^ population while known markers of corticofugal projection neurons (in green) were enriched in the BCL11B^+^ population. Scale bars; 100 *µ*m (b, right panels, d), 500 *µ*m (b, left panel).

To determine whether we isolated the correct neuronal populations and to test nuclear transcriptional profiling using these samples, SMART-Seq v.4 RNA-seq libraries were generated and sequenced on HiSeq 2500. For each sample, libraries were sequenced to a mean of 40 million 100bp paired-end reads (range: 36-48 million reads per sample) to be able to reliably detect low-abundance transcripts. To determine the degree of RNA degradation, we measured the 3’ bias using Qualimap (Okonechnikov et al., 2016). The 3’ bias for P30 samples ranged from 0.65 to 0.69 (mean±SD: 0.685±0.02), which is comparable to RNA Integrity Number (RIN) of 2-4 (Sigurgeirsson et al., 2014) (Figure 1e). Consistent with the idea that nuclear transcripts are predominantly nascent RNA, we found that a substantial number of reads mapped to intronic regions (Exonic: 63.16±4.89%; Intronic: 32.49±4.48%; Intergenic: 4.36±0.56%), (Figure 1f) (Habib et al., 2017; Lake et al., 2016; Lake et al., 2017). Distribution of normalized read counts was virtually identical among samples **(Figure 1-figure supplement 1a)**. Unbiased hierarchical clustering showed that the samples of the same population clustered together (average Pearson correlation between samples within population: r = 0.98) **(Figure 1-figure supplement 1b)**. Subsequently, the two populations were analyzed for differential (gene) expression (DE). The frequency distribution of all *p*-values showed an even distribution of null *p*-values, thus allowing for calculation of adjusted *p*-value using the Benjamini-Hochberg procedure **(Figure 1-figure supplement 1c)**. Between populations, we found 2,698 differentially expressed genes (adjusted *p*-value *<* 0.05) out of 17,662 genes. The high number of genes detected suggests identification of low abundance transcripts.

From the DE analysis, we found an enrichment of known CPN and Layer 4 (L4) markers (e.g. *Cux2*, *Unc5d*, and *Rorb*) in the SATB2^+^ population among the unbiased top 50 DE genes. Conversely, we found an enrichment of CFuPN markers (e.g. *Fezf2*, *Foxp2*, and *Crym*) in the BCL11B^+^ population (Figure 1g). BCL11B also labels interneurons in all layers of the mouse neocortex (Arlotta et al., 2005; Nikouei et al., 2016). Accordingly, we found an enrichment of some interneuron markers in the BCL11B^+^ population (e.g. *Gad1* and *Gad2*) (Figure 1g). To confirm the molecular identities of the isolated neuronal populations, we also determined the relative expression levels of known CPN and CFuPN marker genes that were differentially expressed between CPN and CFuPN in previous studies (21 CPN markers and 22 CFuPN markers) (Arlotta et al., 2005; Molyneaux et al., 2007; Molyneaux et al., 2015). We found that all CPN markers were enriched in the SATB2^+^ population and all CFuPN markers were enriched in the BCL11B^+^ population **(Figure 1-figure supplement 2)**. To validate the differentially expressed genes, we chose four DE genes (*Ddit4l*, *Unc5d*, *Kcnn2*, and *Rprm*) for further analysis. Using RNAscope double fluorescent *in situ* hybridization (FISH), we localized the transcripts of these genes in specific neuronal populations. We found that *Ddit4l* and *Unc5d* were expressed in layers 2 through 4 and were localized to SATB2^+^ neurons **(Figure 1-figure supplement 3)**. Additionally, *Kcnn2* and *Rprm* were expressed in layers 5 and 6, respectively, and they were specifically confined to BCL11B^+^ neurons **(Figure 1-figure supplement 3)**. In addition, we successfully isolated and profiled the same neuronal populations from mature, adult (1+ years old) mouse neocortex **(Figure 1-figure supplement 4-5)**. These results indicate that FIN-Seq can be used to isolate CFuPN and CPN nuclei from flash-frozen mouse neocortex for downstream quantitative RNA-seq analysis of specific neuronal populations.

To determine the degree to which nuclear transcript abundance correlates to cellular transcript abundance, we sought to compare the transcriptional profiles of BCL11B^+^ nuclei and cells. For cells, we dissociated the brains of P7 mice using a protocol described previously (Molyneaux et al., 2015). For nuclei, we performed the FIN-Seq protocol, starting with a fresh P7 brain instead of flash-freezing to keep the starting material consistent between cells and nuclei. We chose the P7 time point because dissociation of the adult mouse brain into single cells affects cell viability at later ages. Transcriptional analysis of BCL11B^+^ cells and nuclei showed a high degree of correlation (average Pearson correlation between cellular vs. nuclear: r = 0.90; cellular vs. cellular: r=0.93; nuclear vs. nuclear: r = 0.93) **(Figure 1-figure supplement 6)**. In contrast, previous comparison of single nucleus and single cell transcriptomes from the adult mouse brain showed a lower degree of correlation (r = 0.77) (Lake et al., 2017). These results indicate that bulk sequencing of isolated nuclei using FIN-Seq could more accurately represent the transcript abundance found within whole cells.

### Specific neuronal subtypes can be isolated from frozen post-mortem human brain samples

To determine whether this protocol is applicable to frozen post-mortem samples of the human brain, we obtained five frozen postmortem brain samples (Brodmann Area 4, primary motor cortex; Ages: 47-61) from a tissue bank that had stored them long-term (for description of the samples, see Materials and Methods). Of note, the oldest frozen sample had been archived for over 25 years. We implemented the same FIN-Seq protocol as above to the frozen human cortical tissue (Figure 2a), in which we found BCL11B^+^/SATB2^−^, BCL11B^+^/SATB2^+^, and BCL11B^−^/SATB2^+^ nuclei (Figure 2b). We used FACS to isolate SATB2^LO^BCL11B^HI^ and SATB2^HI^BCL11B^LO^ nuclei as well as all cortical nuclei (henceforth called BCL11B^+^, SATB2^+^, and All, respectively) for comparison (BCL11B^+^: 26,616 nuclei/replicate, n=5; SATB2^+^: 104,865 nuclei/replicate, n=5; All: 67,580 nuclei/replicate, n=5). These results indicate that nuclear isolation of specific neuronal subtypes from frozen postmortem human brain tissue is feasible using this technique.

**Figure 2:**
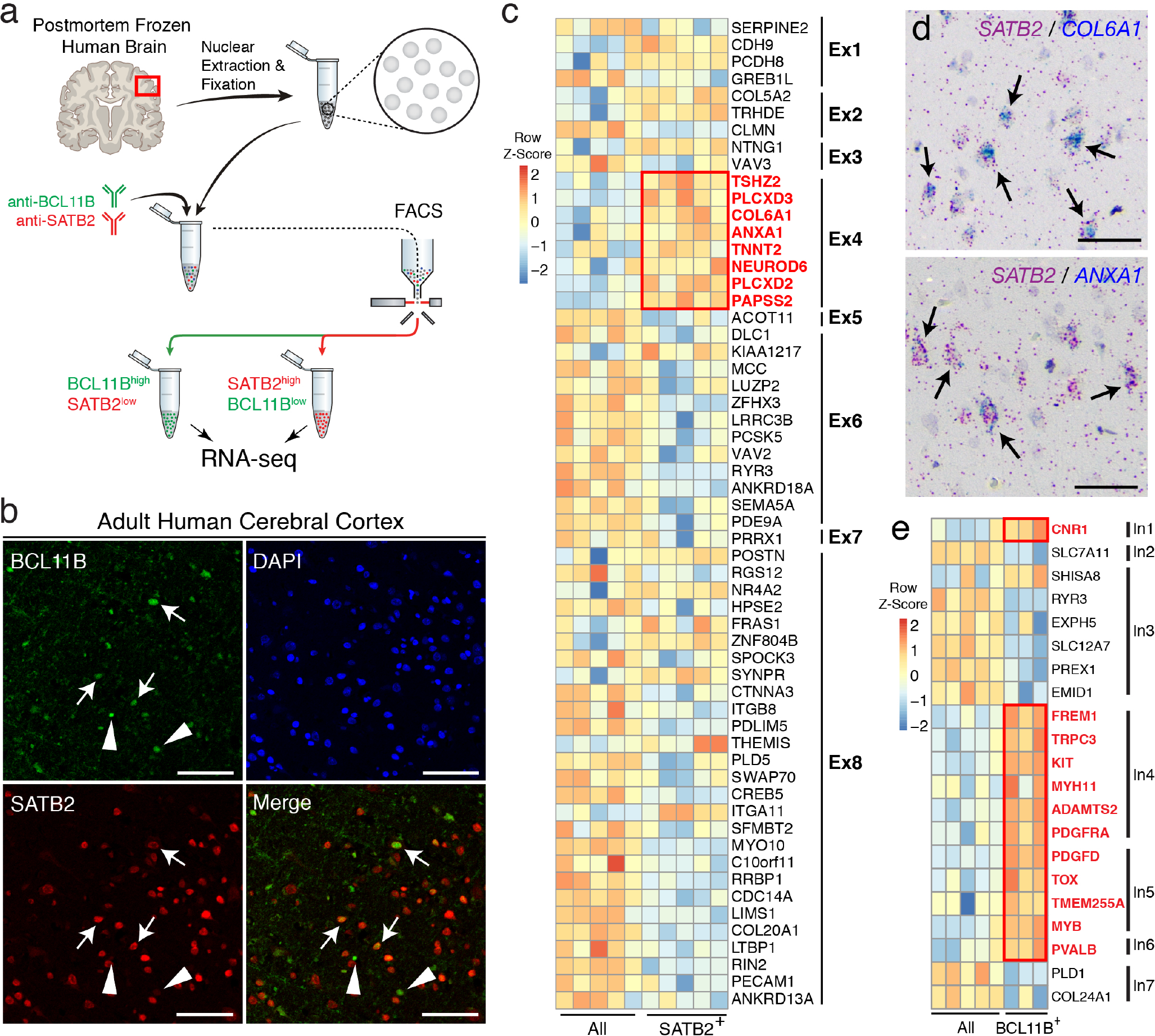
Isolation and profiling of neuronal subtypes from the frozen human cerebral cortex. (**A**) Schematic of FIN-Seq for frozen human cerebral cortex. Nuclei were isolated and subsequently fixed in 4% PFA. They were immunolabeled with anti-BCL11B and anti-SATB2 antibodies, and FACS isolated into populations. RNA from the nuclei were sequenced to obtain a cell type specific transcriptome. (**B**) Representative immunohistochemistry of the adult human cerebral cortex using anti-BCL11B and anti-SATB2 antibodies. Some nuclei expressed both SATB2 and BCL11B (arrows), some nuclei expressed BCL11B but not SATB2 (arrowheads), and many nuclei expressed SATB2 but not BCL11B. (**C**) A heatmap representing relative expression levels of excitatory neuron markers previously identified by single nuclei RNA sequencing that are differentially expressed (adjusted *p*-value*<*0.05) between SATB2^+^ and All populations. Markers of neuronal subtype Ex4 (outlined in red), which expresses SATB2, were enriched in the SATB2^+^ population. (**D**) Validation of Ex4 markers, *COL6A1* (left panel) and *ANXA1* (right panel) using RNAscope single molecule FISH. Both *COL6A1* and *ANXA1* were expressed in SATB2^+^ neurons (arrows). (**E**) A heatmap representing relative expression levels of inhibitory neuron markers previously identified by single nuclei RNA sequencing that are differentially expressed (adjusted *p*-value*<*0.05) between BCL11B^+^ and All populations. Markers of neuronal subtypes, In1, In4, In5, and In6, all of which express BCL11B, were enriched in the BCL11B^+^ population. Scale bars: 100 *µ*m (b), 50 *µ*m (d).

To identify the molecular identity of the isolated neuronal populations, we performed RNA sequencing of each population (BCL11B^+^, SATB2^+^, and All, sequenced to a mean of 36 million paired-end 100bp reads). The average RIN of the frozen human brain samples prior to FIN-Seq was 3.9. After FIN-Seq, the 3’ bias ranged from 0.69 to 0.78 (mean±SD: 0.73±0.02), which corresponds to a RIN of 2-4, indicating that the FIN-Seq protocol does not further decrease the integrity of the RNA **(Figure 2-figure supplement 1a)**. The human brain contains an increased number of nascent transcripts compared to other organs and organisms (Ameur et al., 2011). Accordingly, we found that the proportion of intronic reads was higher in the human neuronal samples compared to that in mice (Exonic: 47.76±5.82%; Intronic: 45.51±5.04%; Intergenic: 6.72±1.13%) **(Figure 2-figure supplement 1b)**. Quality control of the sequencing reads and differential expression analysis indicated successful sample separation and differential expression analysis **(Figure 2-figure supplement 2)**. Between SATB2^+^ and All populations, we found 4,917 differentially expressed genes (adjusted *p*-value *<* 0.05) out of 24,979 genes. Between BCL11B^+^ and All populations, we found 2,812 differentially expressed genes (adjusted *p*-value *<* 0.05) out of 24,477 genes.

To determine the molecular identity of the SATB2^+^ and BCL11B^+^ populations, we first compared the gene expression levels of known markers of oligodendrocytes, astrocytes, and neurons. We found that neuronal markers were enriched in both SATB2^+^ and BCL11B^+^ populations. We also found an enrichment in the BCL11B^+^ population of *PDGFRA*, normally considered an oligodendrocyte marker, but also previously shown to be expressed by a subset of inhibitory neurons in the human cerebral cortex **(Figure 2-figure supplement 3)** (Lake et al., 2016). The SATB2^+^ population highly expressed *SLC17A7* (also known as *VGLUT1*) and did not express *GAD1* or *GAD2*, while the BCL11B^+^ population expressed *GAD1* and *GAD2* at high levels, indicating that, while SATB2^+^ population contained mainly excitatory neurons, BCL11B^+^ population contained also inhibitory neurons **(Figure 2-figure supplement 4)**.

We next sought to understand the identity of the SATB2^+^ and BCL11B^+^ populations at the neuronal subtype-level. Previously, single nucleus RNA-seq has identified eight excitatory neuronal subtypes (Ex1-Ex8) and eight inhibitory neuronal subtypes (In1-In8) in the adult human neocortex (Lake et al., 2016). SATB2 is expressed in all excitatory neurons, but it is most highly expressed in one of the neuronal subtypes referred to, in this prior study, as Ex4. BCL11B is highly expressed in In1, In4, In5, and In6. SATB2 and BCL11B are both expressed in Ex6 and Ex8, but we would not expect to see these subtypes in our populations as we did not collect the SATB2^HI^BCL11B^HI^ population. For the SATB2^+^ population, we cross-referenced our DE gene set (adjusted *p*-value *<* 0.05) to the molecular signature genes that define the eight excitatory cortical neuronal subtypes (Ex1-Ex8). From this analysis, we observed a high level of expression of Ex4 markers in the SATB2^+^ population compared to the All population (Figure 2c). To confirm these results, we also ran the dataset through a gene set enrichment analysis (GSEA) against all marker genes that define Ex1-Ex8 (Subramanian et al., 2005). We found that Ex4 gene set was significantly enriched in the SATB2^+^ population while Ex6 and Ex8 gene sets were enriched in the All population (default significance at FDR *<* 0.25; Ex4: FDR = 0.139; Ex6: FDR = 0.043; Ex8: FDR = 0.005). Depletion of Ex6 and Ex8 from the SATB2^+^ population is likely due to the exclusion of SATB2^HI^BCL11B^HI^ nuclei. We confirmed the expression of *COL6A1* and *ANXA1*, two Ex4 markers, in SATB2^+^ neurons by single molecule FISH (Figure 2d). In the BCL11B^+^ population, we found that the markers for In1, In4, In5, and In6 were enriched compared to the All population (Figure 2e). Furthermore, previous single cell sequencing of the fresh adult human brain identified seven neuronal communities (NC), of which SATB2 is highly expressed in neuronal community 4 (NC4) (Darmanis et al., 2015). Accordingly, we found that the markers for NC4 are highly expressed in the isolated SATB2^+^ population **(Figure 2-figure supplement 5)**. By GSEA analysis, we also found that NC4 gene set was significantly enriched in the SATB2^+^ population (FDR = 0.037). Taken together, our results show the FIN-Seq protocol can isolate molecularly-defined neuronal subtypes for downstream transcriptional profiling from frozen postmortem human cortical samples.

### Isolation and transcriptional profiling of cone photoreceptors from the human retina

To determine whether we could use FIN-Seq to isolate and profile specific cell types from another region of the human CNS, we chose to isolate cone photoreceptors from the retina. We obtained four frozen postmortem eyes (age range: 40-60, see Materials and methods for description of samples) from patients without known retinal disorders. Nuclei were extracted from the mid-peripheral retina, fixed, and immunostained by a human Cone Arrestin (CAR, also known as ARR3) antibody (Figure 3a). In human retinal cross-sections, we found CAR expression in the nuclei and cell bodies of cone photoreceptors, located in the outer nuclear layer where all photoreceptors reside (Figure 3b). CAR^+^ and CAR^−^ nuclei were isolated by FACS, and the RNA was extracted for deep sequencing (CAR^+^: 8,500 nuclei/replicate, n=4; CAR^−^: 180,000 nuclei/replicate, n=4). On average, 1.97% of all nuclei were CAR^+^, a proportion similar to known percentage of cone photoreceptors in the mouse retina (Carter-Dawson and LaVail, 1979). To determine whether fixation was necessary for antibody penetration, we performed the FIN-Seq protocol with and without fixation. We found that the distinct CAR^+^ population was present only with fixation, suggesting that, unlike the NeuN antibody, fixation is necessary for optimal immunolabeling of CAR **(Figure 3-figure supplement 1)**. cDNA sequencing libraries were generated using SMART-Seq v.4 and sequenced to a mean depth of 43 million (range: 37 53 million reads/replicate) 75bp paired-end reads. The sequencing reads were analyzed, and the quality control parameters indicated successful sample separation and differential expression analysis **(Figure 3-figure supplement 2)**. We found 5,260 DE genes (adjusted *p*-value *<* 0.05) between CAR^+^ and CAR^−^ nuclear populations.

**Figure 3:**
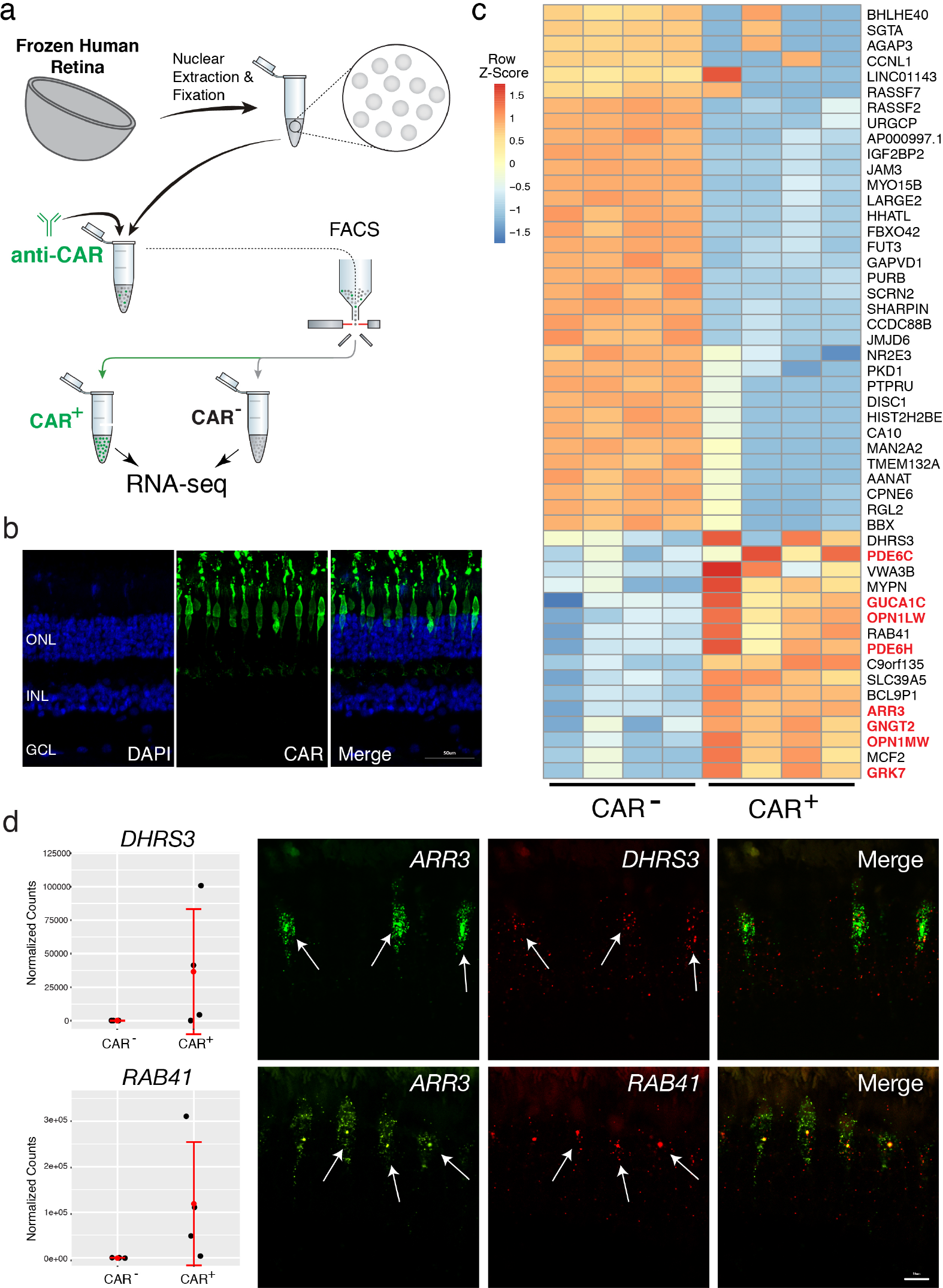
Isolation and sequencing of cone photoreceptor nuclei from the frozen human retina. (**A**) Schematic of FIN-Seq for the frozen human retina. Nuclei were extracted from the frozen retina and subsequently fixed in 4% PFA. Nuclei were then immunolabeled with an anti-CAR antibody and sorted. CAR^+^ and CAR^−^ populations were obtained and the nuclear RNA was sequenced. (**B**) Representative immunohistochemistry of an adult human retina section using the anti-CAR antibody (middle panel) and DAPI (left panel). CAR^+^ cone photoreceptors were localized to the uppermost layer of the ONL. (**C**) Heatmap of unbiased top 50 differentially expressed genes between CAR^+^ and CAR^−^ populations. Known cone photoreceptor markers (in red) were enriched in the CAR^+^ population. (**D**) Validation of new human cone photoreceptor markers by single molecule FISH. Expression levels for *DHRS3* and *RAB41* from the RNA-seq are indicated in the graphs (left panels). Both *DHRS3* and *RAB41* were expressed in the *ARR3*^+^ cone photoreceptors (arrows). ONL, Outer Nuclear Layer; INL, Inner Nuclear Layer; GCL, Ganglion Cell Layer. Scale bars; 50 *µ*m (b), 10 *µ*m (d).

To determine the cellular identity of the CAR^+^ population, we examined the top 50 differential expressed genes between CAR^+^ and CAR^−^ populations. Of the 16 genes enriched in the CAR^+^ population, eight are known markers of human cone photoreceptors, identified by previous single cell RNA sequencing experiments (Figure 3c, **cone markers in red)** (Lukowski et al., 2018). We also performed single molecule FISH for two previously uncharacterized cone markers, *RAB41* and *DHRS3*, and found that they were specifically expressed in ARR3^+^ cone photoreceptors (Figure 3d). To determine whether other cell type specific transcripts were enriched, we assessed the abundance of human markers for cones (*ARR3, GUCA1C, OPN1MW, PDE6C, PDE6H*), rods (*GNGT1, CNGA1*), bipolar cells (*VSX2, TRPM1, GRM6*), amacrine cells (*GAD1*), astrocytes / Müller glia (*GFAP*), Müller glia (*APOE, RLBP1*), and retinal ganglion cells (*SNCG, NEFL*) **(Figure 3-figure supplement 3)**. We found an enrichment of human cone markers in the CAR^+^ population while all other cell type markers were enriched in the CAR^−^ population. These results indicate that FIN-Seq successfully isolated and transcriptionally profiled cone photoreceptors from frozen postmortem human retinas.

## DISCUSSION

Technologies to enable transcriptional analysis of the human CNS are rapidly expanding. At the tissue-level, distinct regions of the fetal and adult human brain have been sampled for gene expression analysis (Bossers et al., 2009; Dangond et al., 2004; Dumitriu et al., 2012; Hauser et al., 2005; Hawrylycz et al., 2012; Kang et al., 2011; Lederer et al., 2007; Miller et al., 2006; Moran et al., 2006; Offen et al., 2009; Papapetropoulos et al., 2006; Wang et al., 2006). Despite progress, these tissue-level approaches cannot account for cellular heterogeneity of the human brain, an organ with tremendous cellular diversity. This is important especially as it refers to human CNS disorders, where histological studies have underscored the cell type specific nature of cellular dysfunction and degeneration (Hartong et al., 2006; Mitchell and Borasio, 2007; Sulzer and Surmeier, 2013). To understand the transcriptional changes that accompany cellular dysfunction in different types of CNS disorders, it is critical to isolate and analyze the specific neuronal populations affected. These may be rare cell types within the tissue, further underscoring the need for technologies that allow enrichment of pre-defined cell types.

Recent developments of single cell RNA-seq technology have enabled unbiased sampling of all cell types from a human CNS tissue sample (Cherry et al., 2018; Darmanis et al., 2015; Hodge et al., 2018; Lake et al., 2016; Lake et al., 2018; Liang et al., 2019; Lukowski et al., 2018; Peng et al., 2019; Phillips et al., 2018). However, for some types of studies, it is impractical to assess the gene expression changes in all cell types. If the cell type of interest is known, bulk RNA-seq of isolated neuronal populations is a complementary approach to quantify gene expression more comprehensively in specific cells of interest. Here, we developed a new method, FIN-Seq, to quantify gene expression in isolated neuronal populations from frozen postmortem human CNS tissue. Bulk RNA sequencing can detect low abundant transcripts and rare splice variants, which are often not detected in single cell or single nucleus RNA sequencing (Arzalluz-Luque and Conesa, 2018; Liu and Trapnell, 2016). We also showed that bulk nuclear sequencing could represent more of the whole-cell transcripts compared to single nucleus sequencing. FIN-Seq is a complementary approach to single nucleus sequencing that can isolate and transcriptionally profile user-defined cell types from frozen human CNS tissues. As suggested by the data from cone photoreceptors, which comprise only 2% of retinal cells, it may prove to be especially valuable for deep profiling of rare cell types.

The challenge of applying FIN-Seq for some cell types is the availability of suitable nuclear antibodies. With the rapid progress of single cell sequencing, markers of molecularly distinct human neuronal subtypes are becoming available. For most of these markers, however, no antibody exists. FIN-Seq could greatly benefit from efforts to generate a validated antibody catalog such as the Protein Capture Reagents Program, in which over 700 validated monoclonal antibodies against human transcription factors have been produced (Venkataraman et al., 2018). For molecular markers without an antibody, FIN-Seq could be further developed to isolate specific cell populations using nuclear RNA by FISH techniques such as RNAscope or SABER (Kishi et al., 2018; Klemm et al., 2014). Labeling specific nuclear transcripts of human neuronal nuclei for downstream FACS and transcriptome sequencing will enable FIN-Seq to capture any cell type of interest.

Taken together, FIN-Seq could enable transcriptional profiling of specific, user-defined neuronal subtypes in the postmortem human CNS without a need for genetic labeling. Counting only those from the NIH brain bank, over 16,000 postmortem samples are available, including those with neurological disorders, and many of them are stored long-term as flash-frozen samples. With FIN-Seq, we can start to interrogate the transcriptional changes that accompany specific neuronal subtypes in the adult human brain and identify molecular mechanisms underlying cell type specific pathology.

## MATERIALS AND METHODS

### Mouse Brain Samples

All animals were handled according to protocols approved by the Institutional Animal Care and Use Committee (IACUC) of Harvard University. For each biological replicate, the neocortex of P30 or adult (1+ years old) CD1 mice were microdissected, flash-frozen in an isopentane/dry ice slurry, and stored at −80°C.

### Frozen Human CNS Samples

Frozen Brodmann Area 4 (Primary Motor Cortex) samples of Patient 1569, 3529, 3589, 4340, and 5650 were obtained from Human Brain and Spinal Fluid Resource Center at University of California, Los Angeles through the NIH NeuroBioBank. Patient 1569 is a 61-year-old male with no clinical brain diagnosis and the postmortem interval was 9 hours. Patient 3529 is a 58-year-old male with no clinical brain diagnosis and the postmortem interval was 9 hours. Patient 3589 is a 53-year-old male with no clinical brain diagnosis and the postmortem interval was 15 hours. Patient 4340 is a 47-year-old male with no clinical brain diagnosis and the postmortem interval was 12.5 hours. Patient 5650 is a 55-yearold male with no clinical brain diagnosis and the postmortem interval was 22.6 hours. This IRB protocol (IRB16-2037) was determined to be not human subjects research by the Harvard University-Area Committee on the Use of Human Subjects.

Frozen eyes were obtained from Restore Life USA (Elizabethton, TN) through TissueForResearch. Patient DRLU032618A is a 52-year-old female with no clinical eye diagnosis and the postmortem interval was 8 hours. Patient DRLU041518A is a 57-year-old male with no clinical eye diagnosis and the postmortem interval was 5 hours. Patient DRLU041818C is a 53-year-old female with no clinical eye diagnosis and the postmortem interval was 9 hours. Patient DRLU051918A is a 43-year-old female with no clinical eye diagnosis and the postmortem interval was 5 hours. This IRB protocol (IRB17-1781) was determined to be not human subjects research by the Harvard University-Area Committee on the Use of Human Subjects.

### Nuclei Isolation, Immunolabeling, and FACS

Nuclei were prepared as described previously (Krishnaswami et al., 2016), with modifications. Thawed tissue was minced and incubated in 1% PFA (with 1 *µ*L mL^−1^ RNasin Plus (Promega, Madison, WI)) for 5 minutes. Nuclei were prepared by Dounce homogenizing in 0.1% Triton X-100 homogenization buffer (250 mM sucrose, 25 mM KCl, 5 mM MgCl2, 10mM Tris buffer, pH 8.0, 1 *µ*M DTT, 1× Protease Inhibitor (Promega), Hoechst 33342 10 ng mL^−1^ (Thermo Fisher Scientific, Waltham, MA), 0.1% Triton X-100, 1 *µ*L mL^−1^ RNasin Plus). Sample was then overlaid on top of 20% sucrose bed (25 mM KCl, 5 mM MgCl2, 10mM Tris buffer, pH 8.0) and spun at 500xg for 12 minutes at 4°C. The pellet was resuspended in 4% PFA (with 1 *µ*L mL^−1^ RNasin Plus) and incubated for 15 minutes on ice. The sample was spun at 2000xg for 5 minutes at 4°C and the supernatant was discarded. The sample was then resuspended in blocking buffer (0.5% BSA in nuclease-free PBS, 0.5 *µ*L mL^−1^ RNasin Plus) and incubated for 15 minutes. Sample was spun and the pellet was resuspended and incubated in primary antibody (1:50 SATB2 antibody (Abcam, Cambridge, UK), 1:100 BCL11B antibody (Abcam), 1:1000 CAR antibody (kind gift from Dr. Sheryl Craft) in blocking buffer) for 30 minutes at 4°C. After washing 1× with blocking buffer, the sample was incubated in secondary antibody (1:750 appropriate AlexaFluor secondary antibodies (Thermo Fisher Scientific)) for 30 minutes at 4°C. After 1× wash, the sample was passed through a 35*µ*m filter (Corning, Corning, NY) before proceeding to FACS. 2N nuclei were determined by Hoechst histogram, and isolated populations were sorted into blocking buffer. Sorted nuclei were spun at 3000×g for 7 minutes, and the supernatant was discarded.

### RNA isolation and library preparation

RNA was extracted using the RecoverAll Total Nuclear Isolation Kit (Thermo Fisher Scientific). Crosslinking was reversed by incubating the nuclear pellet in Digestion Buffer and Protease mixture (100 *µ*L buffer and 4 *µ*L protease) for 3 hours at 50°C. RNA-seq library was generated using the SMART-seq v.4 Ultra Low Input RNA Kit (Takara Bio, Kusatsu, Japan) and Nextera XT DNA Library Prep Kit (Illumina, San Diego, CA) according to protocol. Number of cycles was determined based on the number of nuclei sorted. The cDNA library fragment size was determined by BioAnalyzer 2100 HS DNA Assay (Agilent, Santa Clara, CA). The libraries were sequenced as paired-end reads on HiSeq 2500 or NextSeq 500.

### RNA-seq data processing

Quality control of RNA-seq reads were performed using fastqc version 0.11.5 (https://www.bioinformatics.babraham.ac.uk/projects/fastqc/). RNA-seq reads were clipped and mapped onto the mouse genome (Ensembl GRCm38.88) or human genome (Ensembl GRCh38.87) using STAR version 2.5 (Aken et al., 2017; Dobin et al., 2013). Parameters used were as follows: –runThreadN 6 –readFilesCommand zcat –outSAMtype BAM SortedByCoordinate –outSAMunmapped Within –outSAMattributes Standard – clip3pAdapterSeq –quantMode TranscriptomeSAM GeneCounts

Alignment quality control was performed using Qualimap version 2.2.1 (Okonechnikov et al., 2016). Read counts were generated by HT-seq version 0.6.1 (Anders et al., 2015). Sample parameters used were as follows: -i gene name -s no

The resulting matrix of read counts were analyzed for differential expression by DESeq2 version 3.5 (Love et al., 2014). Samples with non-neuronal cell contamination were discarded for analysis (BCL11B^+^ 3529 and BCL11B^+^ 3589). For the DE analysis of human retina samples, any genes with more than 5 samples with zero reads were discarded. The R scripts used for differential expression analysis is available in Supplementary Files.

### Gene Set Enrichment Analysis

GSEA analysis was performed on the All vs. SATB2^+^ dataset using GSEA v3.0 (Subramanian et al., 2005). Gene set databases including markers that define neuronal subtypes identified by Darmanis et al. (2015) and Lake et al. (2016) were generated. Parameters used were as follows: Number of permutations: 1000; Enrichment statistic: weighted; Metric for ranking genes: Signal2Noise; Min size: 0. To determine significance, we used the default FDR *<* 0.25 for all gene sets.

### RNAscope

P30 and adult mouse brains were perfused with 4% PFA and sectioned on a cryostat at a thickness of 16 *µ*m. Double *in situ* fluorescence hybridization was performed using the RNAscope Fluorescent Multiplex assay according to protocol (Advanced Cell Diagnostics, Newark, CA). The following probes were used for the mouse study: Satb2-C1 (Catalog : 413261), Satb2-C2 (Catalog: 420981-C2), Bcl11b-C3 (Catalog: 413271-C3), Ddit4l-C1 (Catalog: 468551), Kcnn2-C1 (Catalog: 427971), Unc5d-C2 (Catalog: 480461-C2), and Rprm-C2 (Catalog: 466071-C2).

FFPE adult human cerebral cortex tissue from a 54-year-old female was obtained from Abcam (ab4296). Chromogenic double *in situ* hybridization was performed for the human brain tissue using the RNAscope 2.5 HD Duplex Assay (Advanced Cell Diagnostics) according to protocol. Fluorescent Multiplex assay was used for the human retina tissue according to protocol (Advanced Cell Diagnostics). The following probes were used for the human study: SATB2-C2 (Catalog: 420981-C2), RORB-C1 (Catalog: 446061), UNC5D-C1 (Catalog: 459991), CRYM-C2 (Catalog: 460031-C2), GAD1-C1 (Catalog: 404031), COL6A1-C1 (Catalog: 482461), ANXA1-C1 (Catalog: 465411), ARR3-C2 (Catalog: 486461-C2), DHRS3-C1 (Catalog: 504941), RAB41-C1 (Catalog: 559561).

### Immunohistochemistry

P30 and adult mouse brains were perfused with 4% PFA and sectioned on a vibratome at a thickness of 40 *µ*m. Immunohistochemistry was performed as previously described (Arlotta et al., 2005) with anti-SATB2 and anti-BCL11B antibodies. FFPE adult human cerebral cortex tissue from a 54-year-old female was obtained from Abcam (ab4296). The brain tissue was deparaffinized by 2× xylene incubation (3 minutes each) followed by 1× 100% ethanol (3 minutes), 1× 95% ethanol (3 minutes), 1× 70% ethanol (3 minutes) washes. Antigen re-trieval was performed in a citrate buffer (10mM Citric Acid, pH 6.0) in a rice cooker with boiling water for 20 minutes. Subsequently, immunohistochemistry was performed as described above with an additional step of incubation in True-Black (Biotium, Fremont, CA) after incubation in blocking buffer to quench the lipofuscin autofluorescence. For human eye immunohistochemistry, formalin-fixed human postmortem eyes were obtained from Restore Life USA. Patient DRLU101818C is a 54-year-old male with no clinical eye diagnosis and the postmortem interval was 4 hours. Patient DRLU110118A is a 59-year-old female with no clinical eye diagnosis and the postmortem interval was 4 hours. The retina was cryosectioned at 16 *µ*m thickness. Immunohistochemistry was performed as previously described (Arlotta et al., 2005) with anti-CAR antibody (1:10,000).

### Imaging

Fluorescent confocal images of the brain were acquired with Zeiss LSM 700 confocal microscope and analyzed with the ZEN Black software. Fluorescent confocal images of the retina were acquired with Nikon Ti inverted spinning disk microscope and analyzed with the NIS-Elements software. Brightfield color images of the human brain were acquired with AxioZoom V16 Zoom Microscope.

## Acknowledgments

We would like to thank former and current members of the Arlotta and Cepko Labs for the insightful discussion and feedback. We thank D. Richardson, Harvard Center for Biological Imaging, P.M. Llopis, R. Stephansky, and the MicRoN core at Harvard Medical School for their assistance with microscopy. We thank S.H. Sui, V. Barrera, and M. Ziller for their assistance with bioinformatics. We thank S. Craft for sharing of the hCAR antibody. We thank C. Araneo, F. Lopez, and the Flow Cytometry Core Facility for their assistance with flow cytometry.

## Author Contributions

R.A., E.Z., H.-H.C., and P.A. conceived the work and designed the experiments. R.A. performed the majority of the experiments and analyzed the data. E.Z. and P.A. conceived the method.E.Z. and H.-H.C. assessed feasibility in preliminary steps. R.A., P.A., and C.L.C. wrote the manuscript. E.Z. edited the manuscript. N.C.C. and H.-H.C. assisted with the FACS. S.K. assisted with the initial antibody screening for the FACS. P.A. and C.L.C. supervised the brain and retina aspects of the project, respectively.

