## Supplementary Figures for "FIN-Seq: Transcriptional profiling of specific cell types in frozen archived tissue from the human central nervous system"

**Figure Supplements**

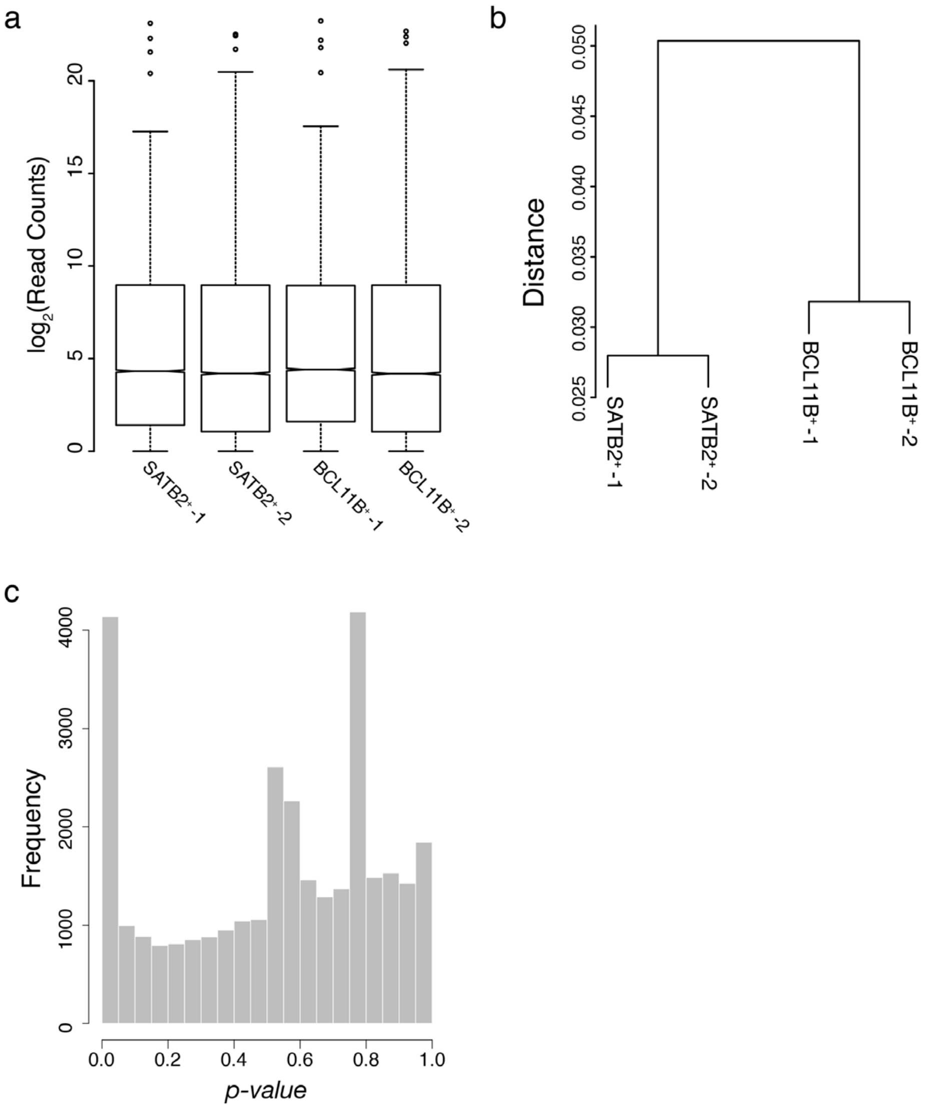


**Figure 1 - figure supplement 1: Quality control for P30 mouse FIN-Seq data.**

a) $Log_2$-transformed read distribution plot for sequenced mouse cortical samples. b) Dendrogram of read counts shows clustering of $SATB2^+$ samples and $BCL11B^+$ samples. c) A plot of frequencies of *p*-values shows an even distribution of null *p*-values.

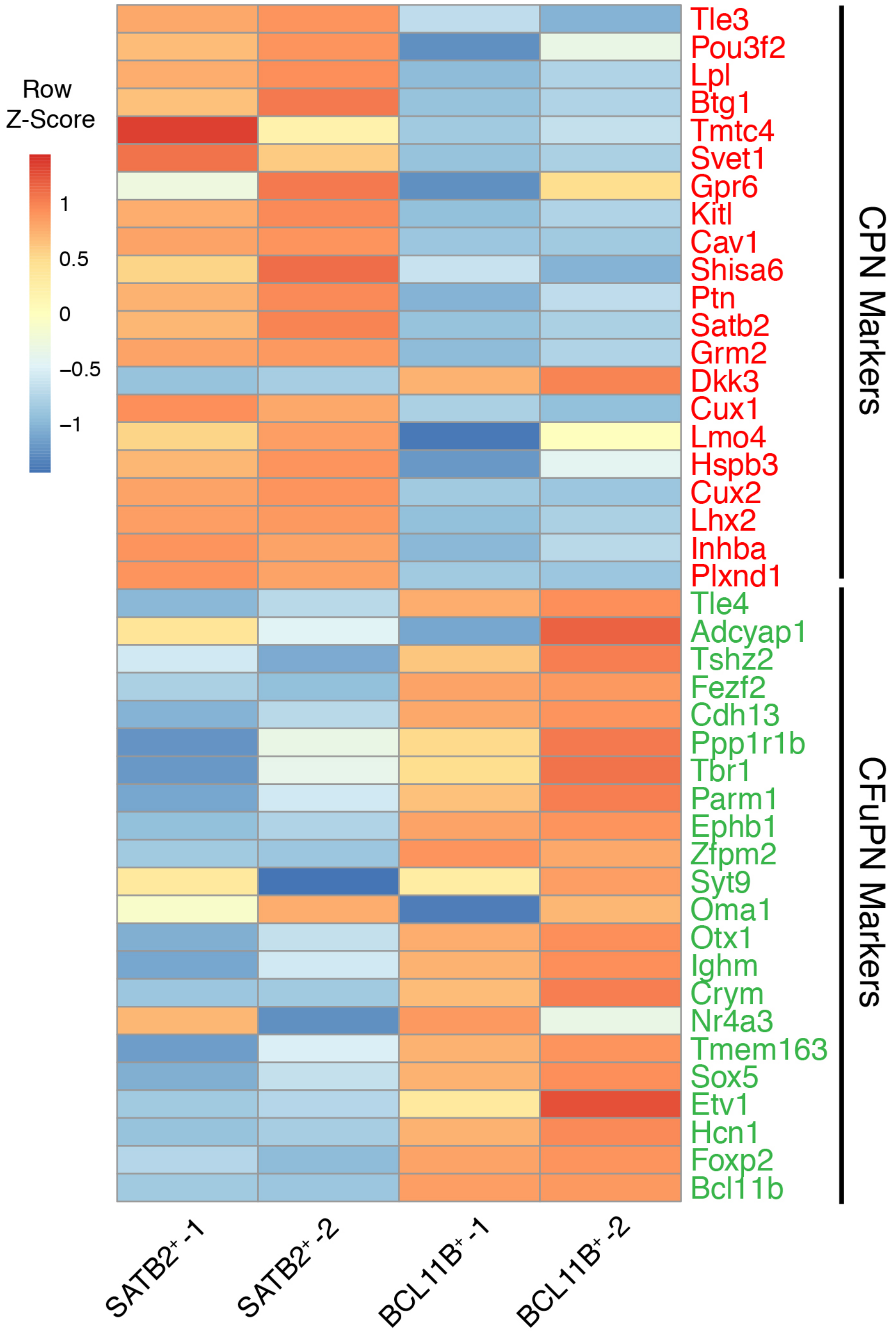


**Figure 1 - figure supplement 2: Heatmap of known CPN and CFuPN markers.**

CPN markers (in red) were all enriched in the SATB2$^{+}$ population while CFuPN markers (in green) were all enriched in the BCL11B$^{+}$ population.

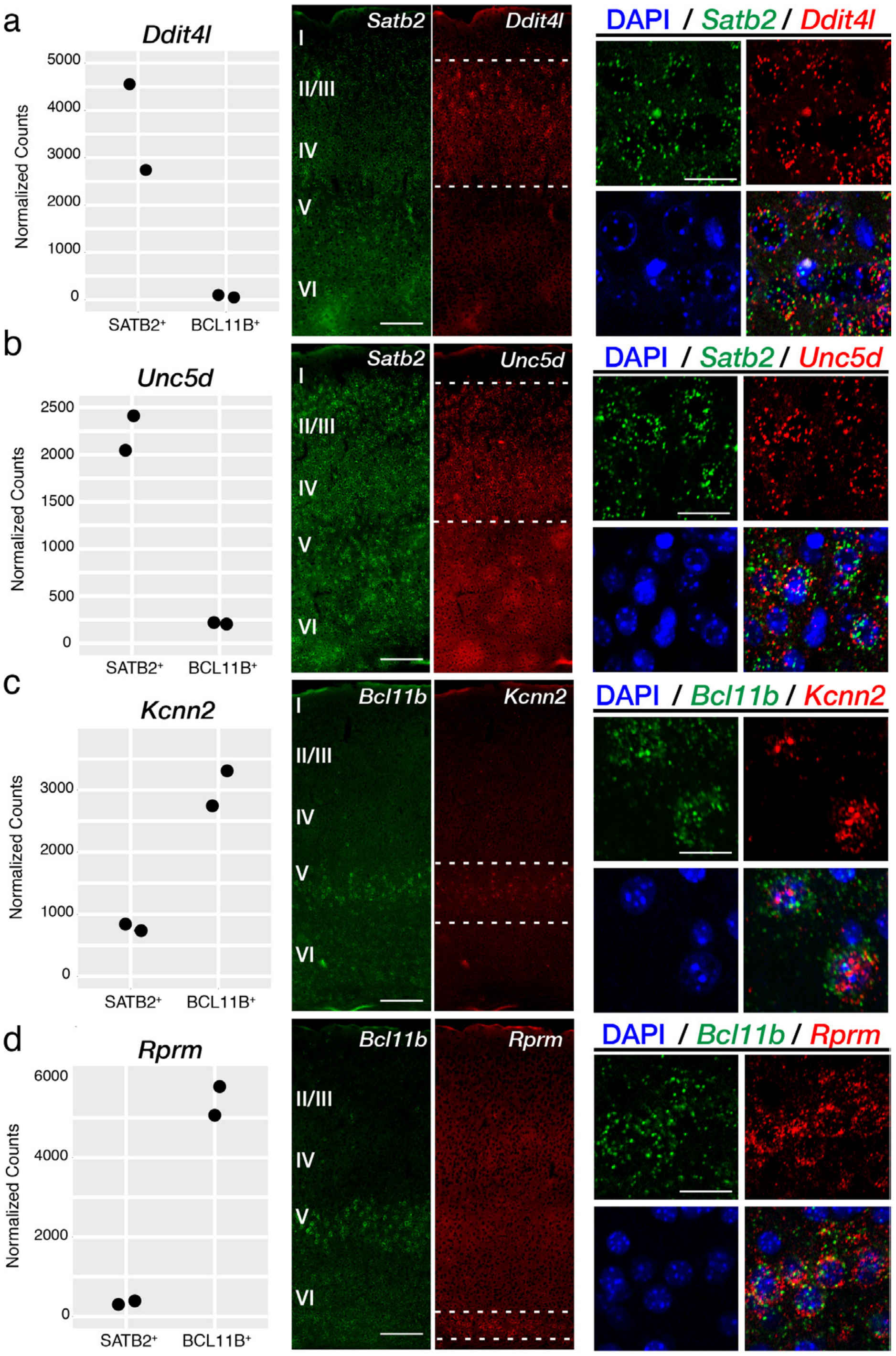


**Figure 1 - figure supplement 3: Validation of subtype-specific transcripts by single molecule FISH.**

Expression values from the RNA-seq are plotted (left panel). *Ddit4l* (a) and *Unc5d* (b) were expressed in the upper layers at P30 in the mouse neocortex (middle panel, between white dotted lines). At this age, *Satb2* was expressed in subset of neurons of all layers (left panel). Merged higher magnification inset (right panel) shows expression of *Ddit4l* (a) and *Unc5d* (b) in *Satb2*$^{+}$ cells in layer 2/3. *Kcnn2* (c) and *Rprm* (d) were expressed in layer 5 and 6, respectively, in the mouse neocortex (middle panel, between white dotted lines). *Bcl11b* was expressed in neurons of layer 5 (left panel). Merged higher magnification inset (right panel) shows expression of *Kcnn2* (c) and *Rprm* (d) in *Bcl11b*$^{+}$ cells. Scale bars; 500 μm (a-d, middle panels), 20 μm (a-d, right panels).

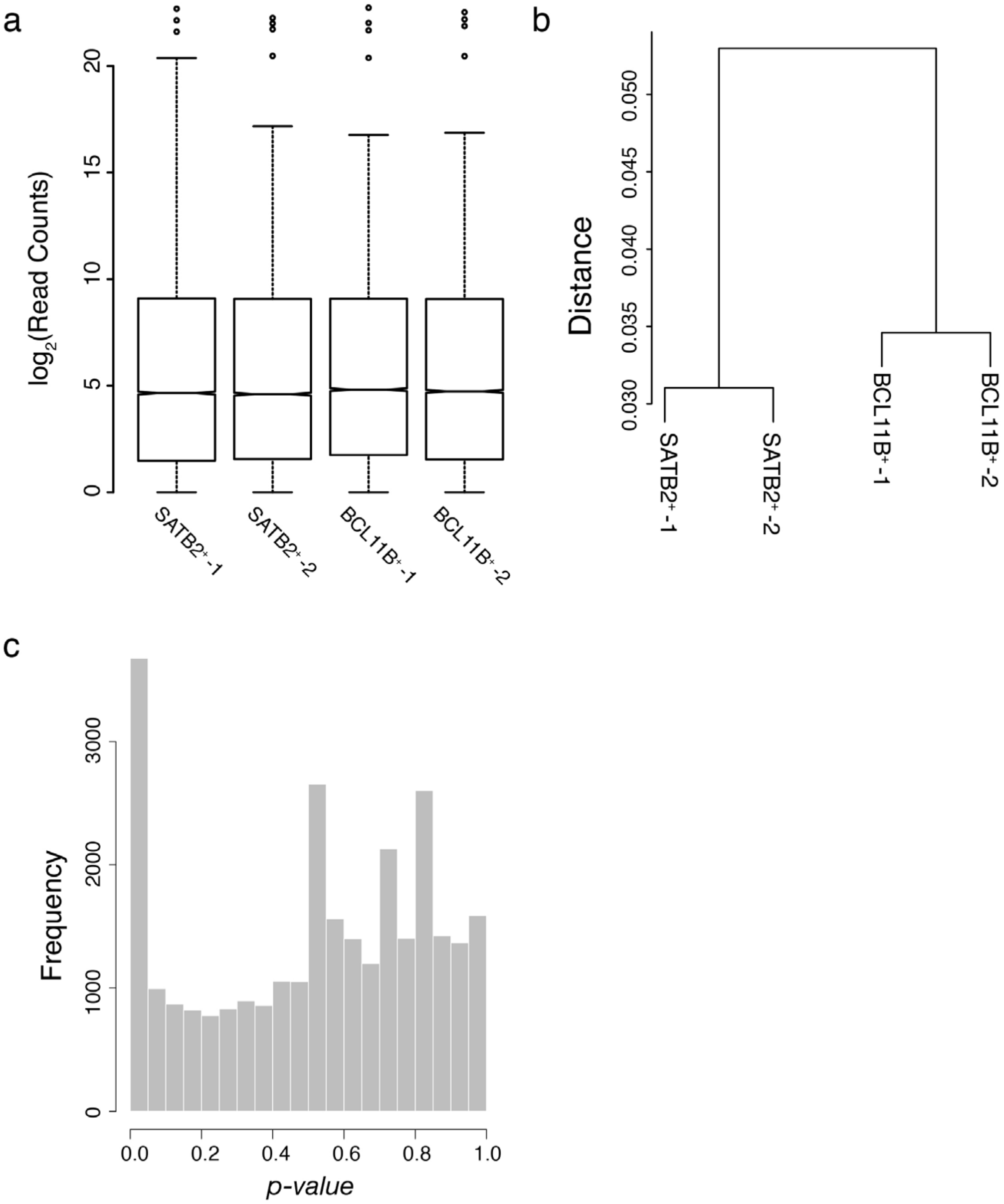


**Figure 1 - figure supplement 4: Quality control for adult mouse FIN-Seq data.**

a) $Log_2$-transformed read distribution plot for sequenced mouse cortical samples. b) Dendrogram of read counts shows clustering of $SATB2^+$ samples and $BCL11B^+$ samples. c) A plot of frequencies of *p*-values shows an even distribution of null *p*-values.

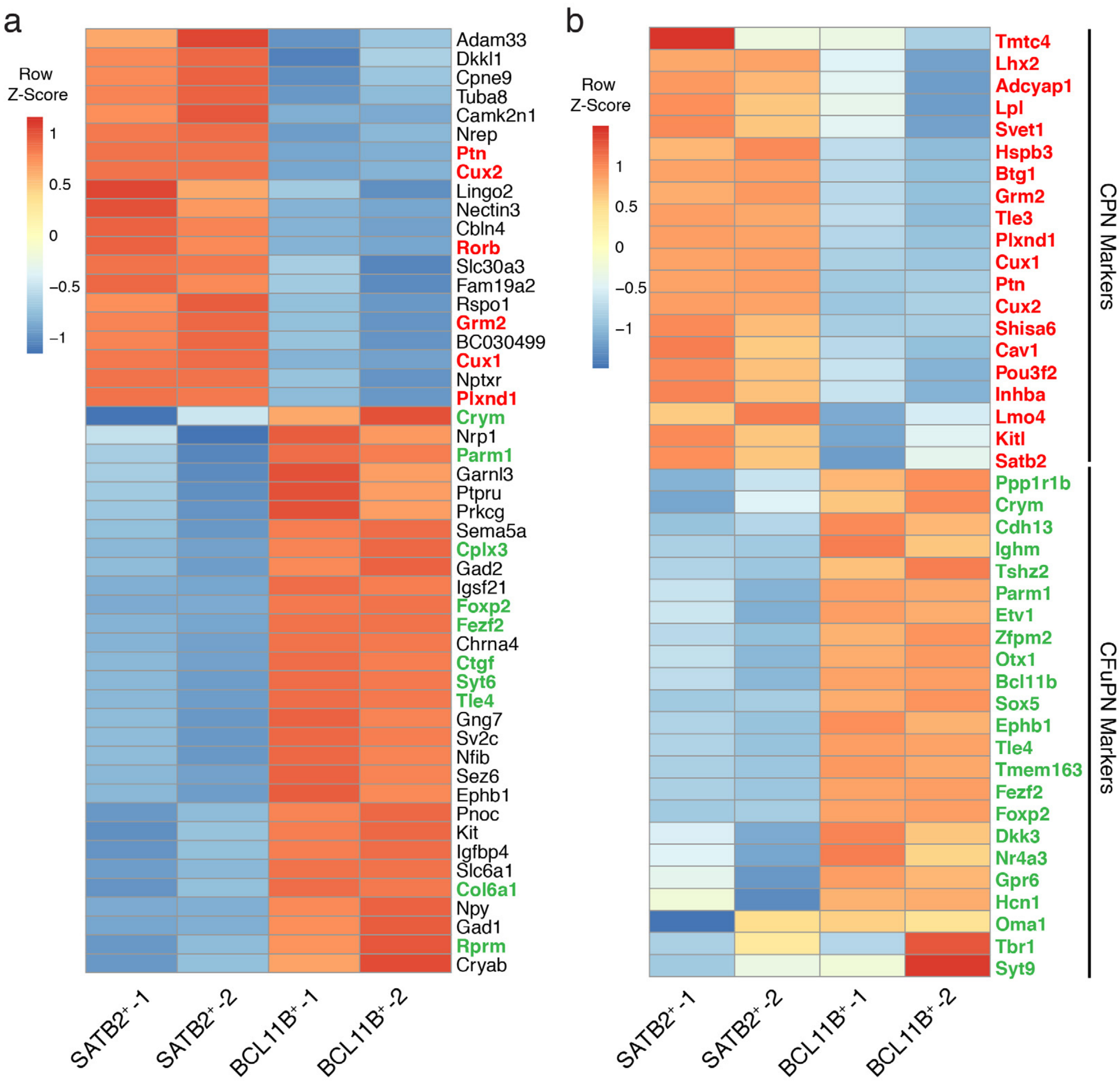


**Figure 1 - figure supplement 5: Heatmap of unbiased top 50 genes and known CPN and CFuPN markers for the adult mouse FIN-Seq data.**

a) Heatmap of unbiased top 50 differentially expressed genes between SATB2$^+$ and BCL11B$^+$ populations. Known markers of CPN (in red) were enriched in the SATB2$^+$ population while known markers of CFuPN (in green) were enriched in the BCL11B$^+$ population. b) CPN markers (in red) were all enriched in the SATB2$^+$ population while CFuPN markers (in green) were all enriched in the BCL11B$^+$ population.

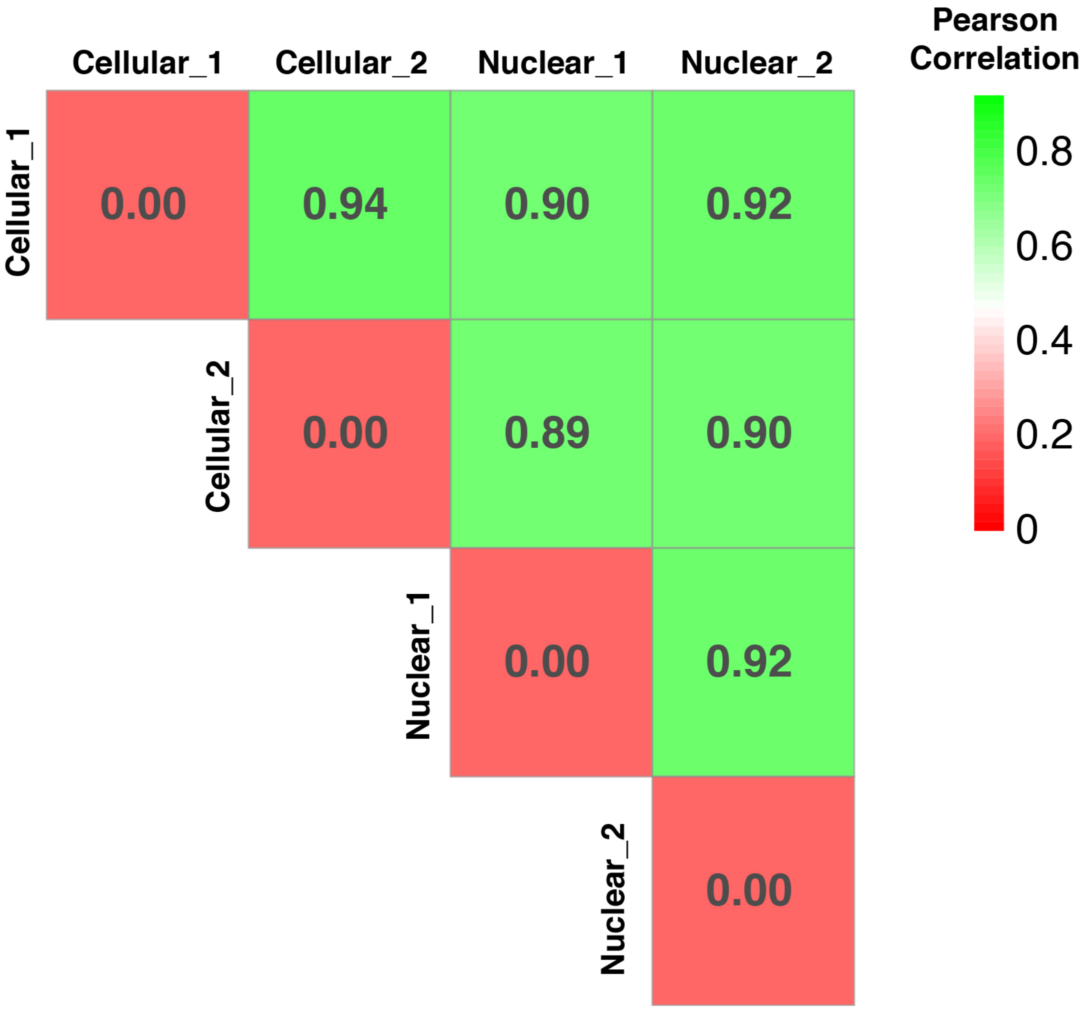


**Figure 1 - figure supplement 6: Heatmap of correlation between cellular and nuclear transcriptomes.**

Pearson correlation between cellular P7 BCL11B$^{+}$ transcriptomes (n=2) and nuclear P7 BCL11B$^{*}$ transcriptomes (n=2) shows high correlation between cellular and nuclear transcriptomes.

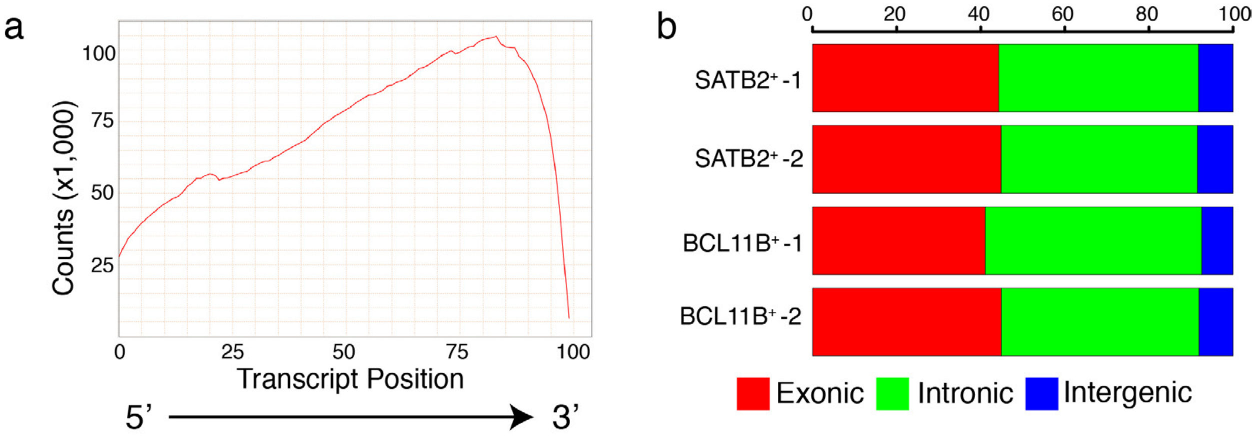


**Figure 2 - figure supplement 1: Quality control for genome mapping of human cortical FIN-Seq reads.**

a) Quantification of read counts mapped to transcript position (5’ to 3’) for every gene shows a strong 3’ bias. b) Quantification of percentage of read counts mapped to exonic, intronic, or intergenic regions of the genome shows high percentage of intronic (green) reads.

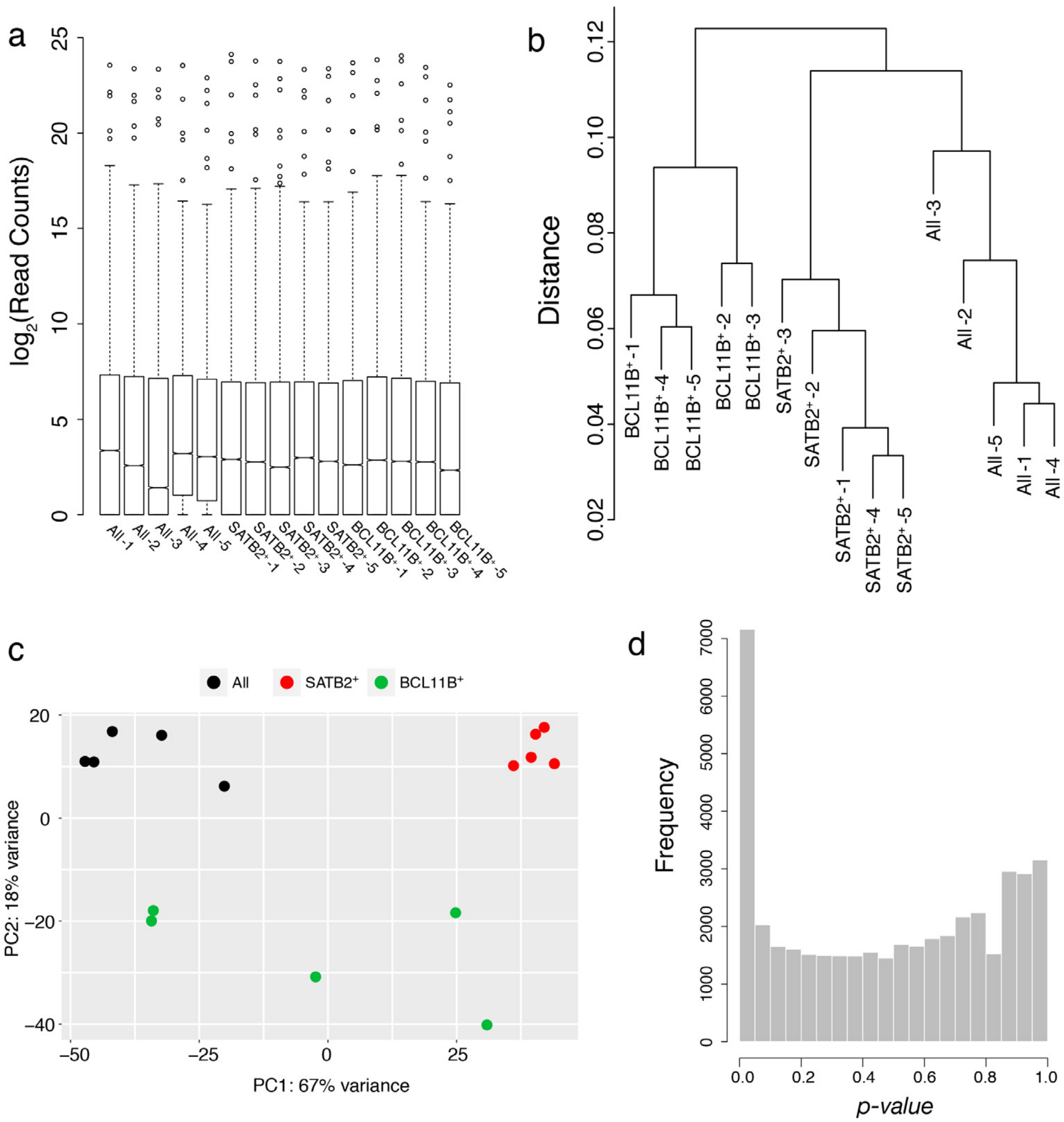


**Figure 2 - figure supplement 2: Quality control for adult human cortical FIN-Seq data.**

a) $Log_2$-transformed read distribution plot for sequenced human cortical samples. b) Dendrogram of read counts shows clustering of SATB2$^+$ samples, BCL11B$^+$ samples, and All samples. c) PCA plot shows clustering of SATB2$^+$ samples, BCL11B$^+$ samples, and All samples. d) A plot of frequencies of *p*-values shows an even distribution of null *p*-values.

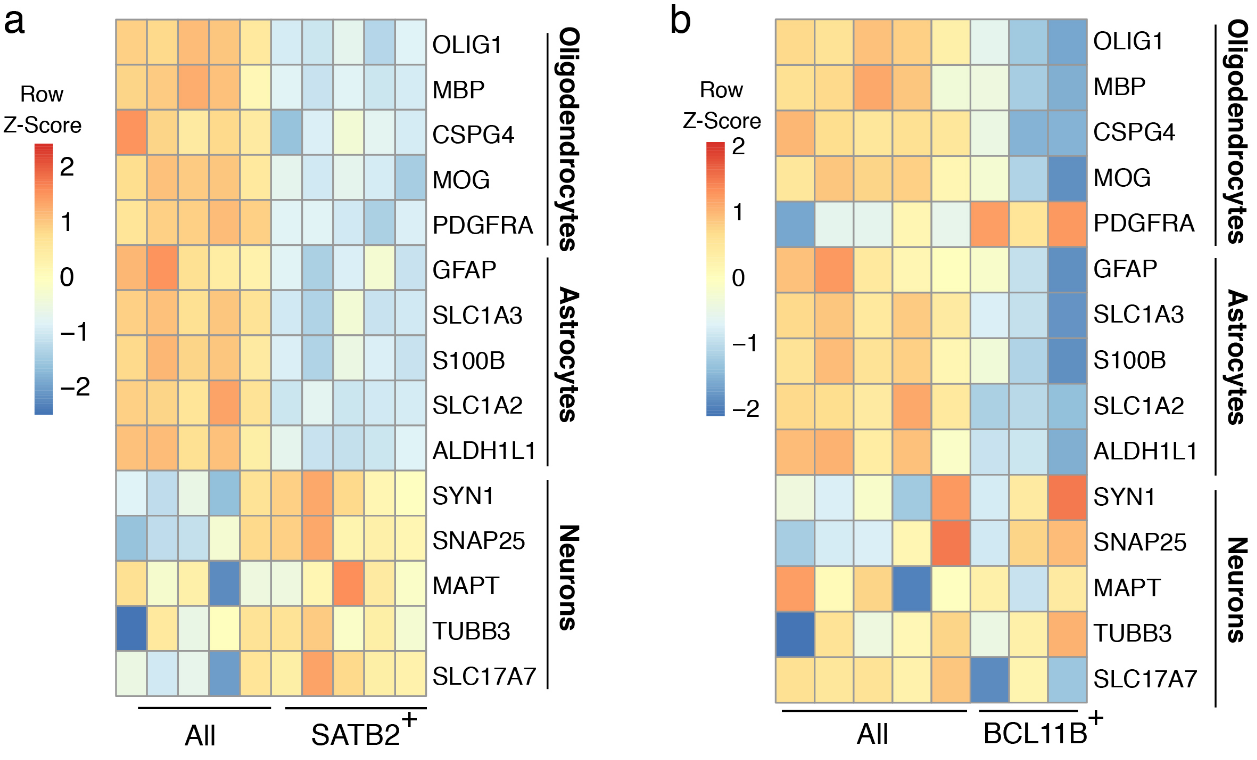


**Figure 2 - figure supplement 3: BCL11B$^+$ population contains specific inhibitory neuronal subtypes.**

A heatmap representing relative expression levels of cell class markers in All and SATB2$^+$ (a) or All and BCL11B$^+$ (b) populations. Expression profile of five oligodendrocyte markers (OLIG1, MBP, CSPG4, MOG, PDGFRA), five astrocyte markers (GFAP, SLC1A3, S100B, SLC1A2, ALDH1L1), and five neuronal markers (SYN1, SNAP25, MAPT, TUBB3, SLC17A7) shows enrichment of neuronal markers in both populations.

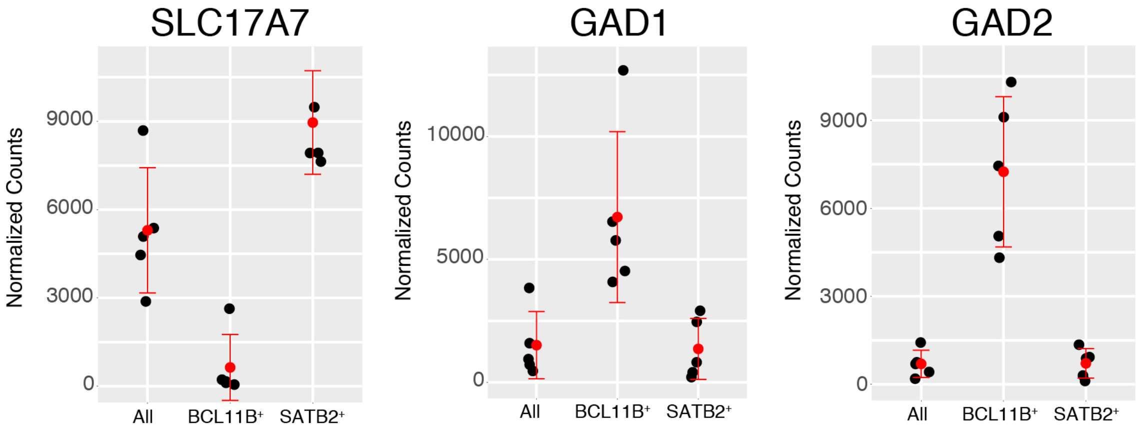


**Figure 2 - figure supplement 4: SATB2$^{+}$ population contains mostly excitatory neurons while BCL11B$^{+}$ neurons also contain inhibitory neurons.**

SLC17A7, an excitatory neuron marker, was enriched in SATB2$^{+}$ neurons (left panel). GAD1 and GAD2, inhibitory neuron markers, were highly expressed in the BCL11B$^{+}$ neurons in the human cerebral cortex (middle and right panels).

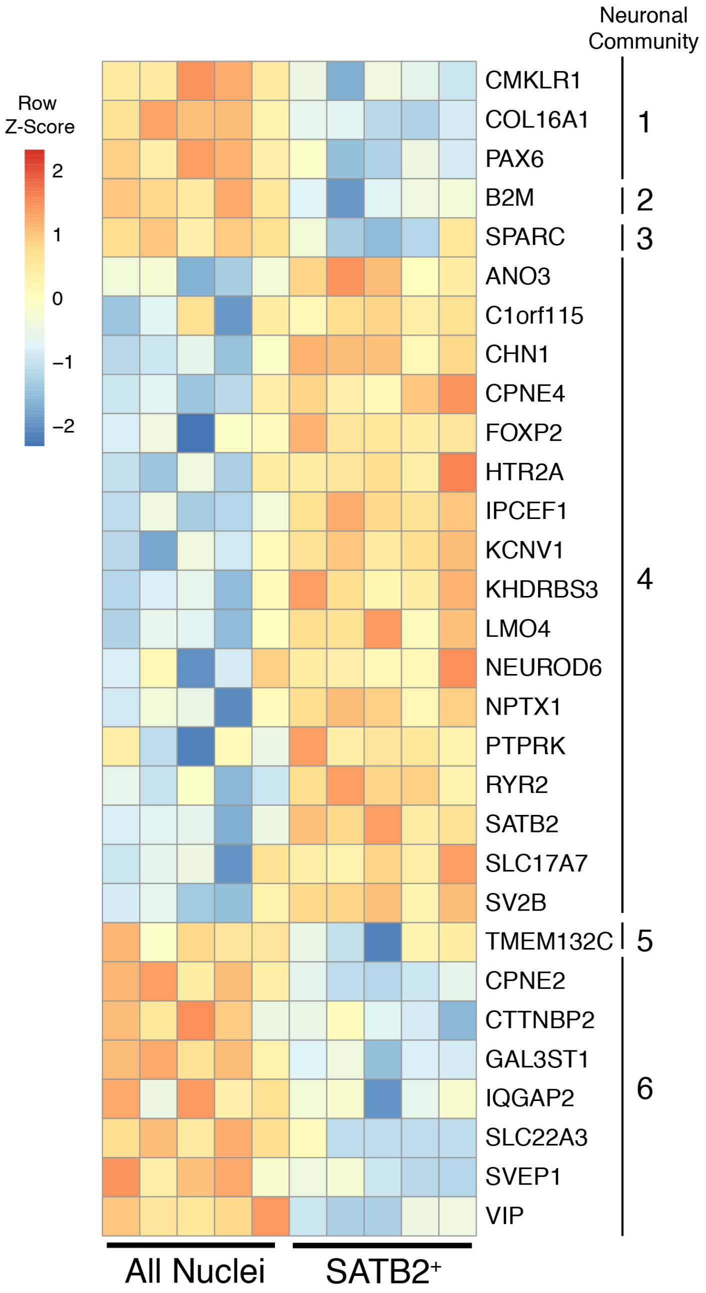


**Figure 2 - figure supplement 5: $SATB2^+$ population represents neuronal community 4.**

A heatmap representing relative expression levels of neuronal community markers previously identified by single cell RNA sequencing of a fresh human cortical sample. Markers of neuronal community 4, which expresses SATB2, were enriched in the $SATB2^+$ population.

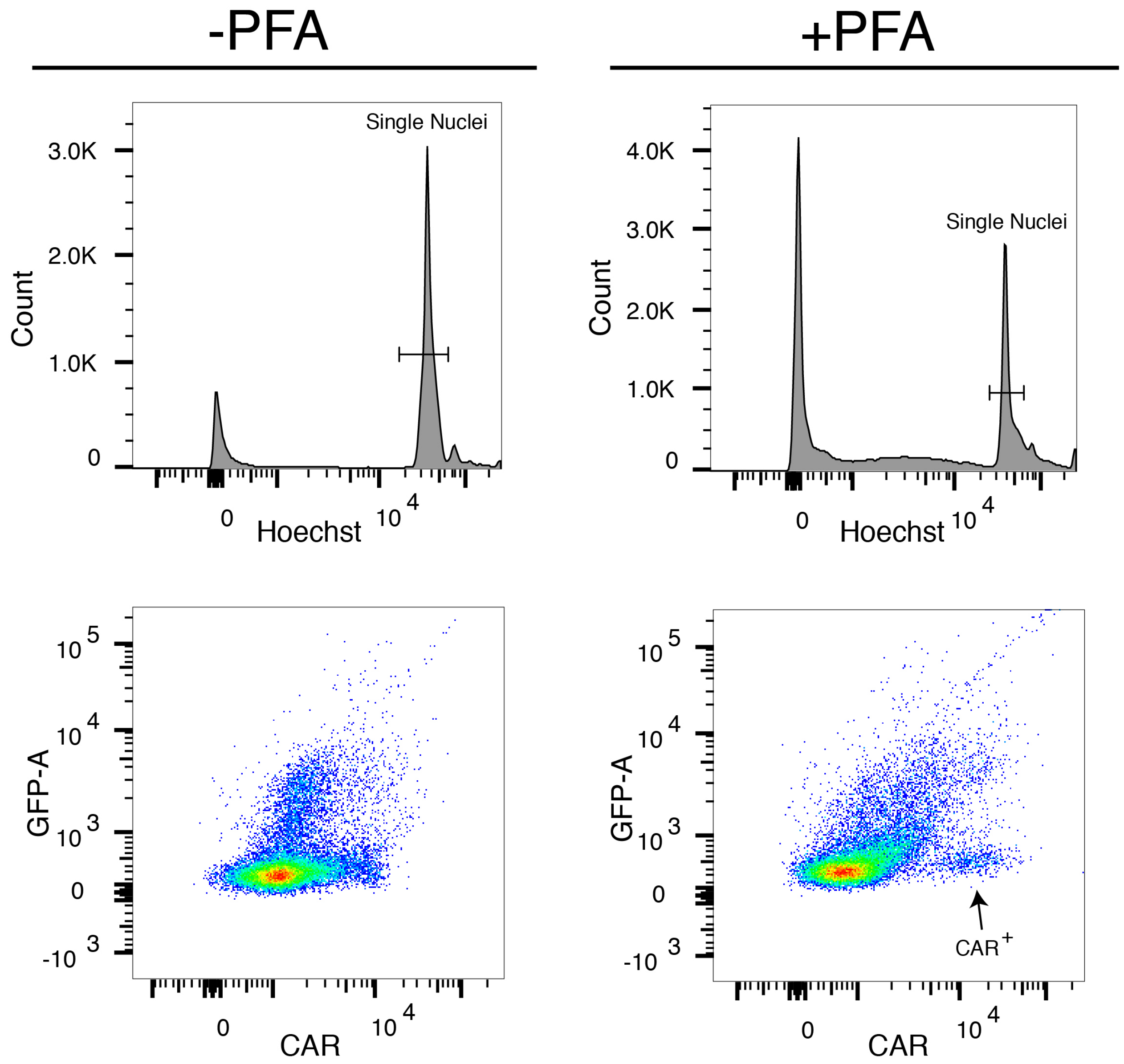


**Figure 3 - figure supplement 1: Fixation is necessary for labeling cone photoreceptor nuclei with CAR.**

FACS plots displaying histogram of Hoechst$^+$ nuclei with and without 4% PFA (upper panels: left panel without PFA, right panel with PFA). FACS plots displaying intensity of CAR staining on the x-axis and 488 nm autofluorescence on the y-axis (bottom panels). CAR$^+$ cluster was only visible with PFA (arrow).

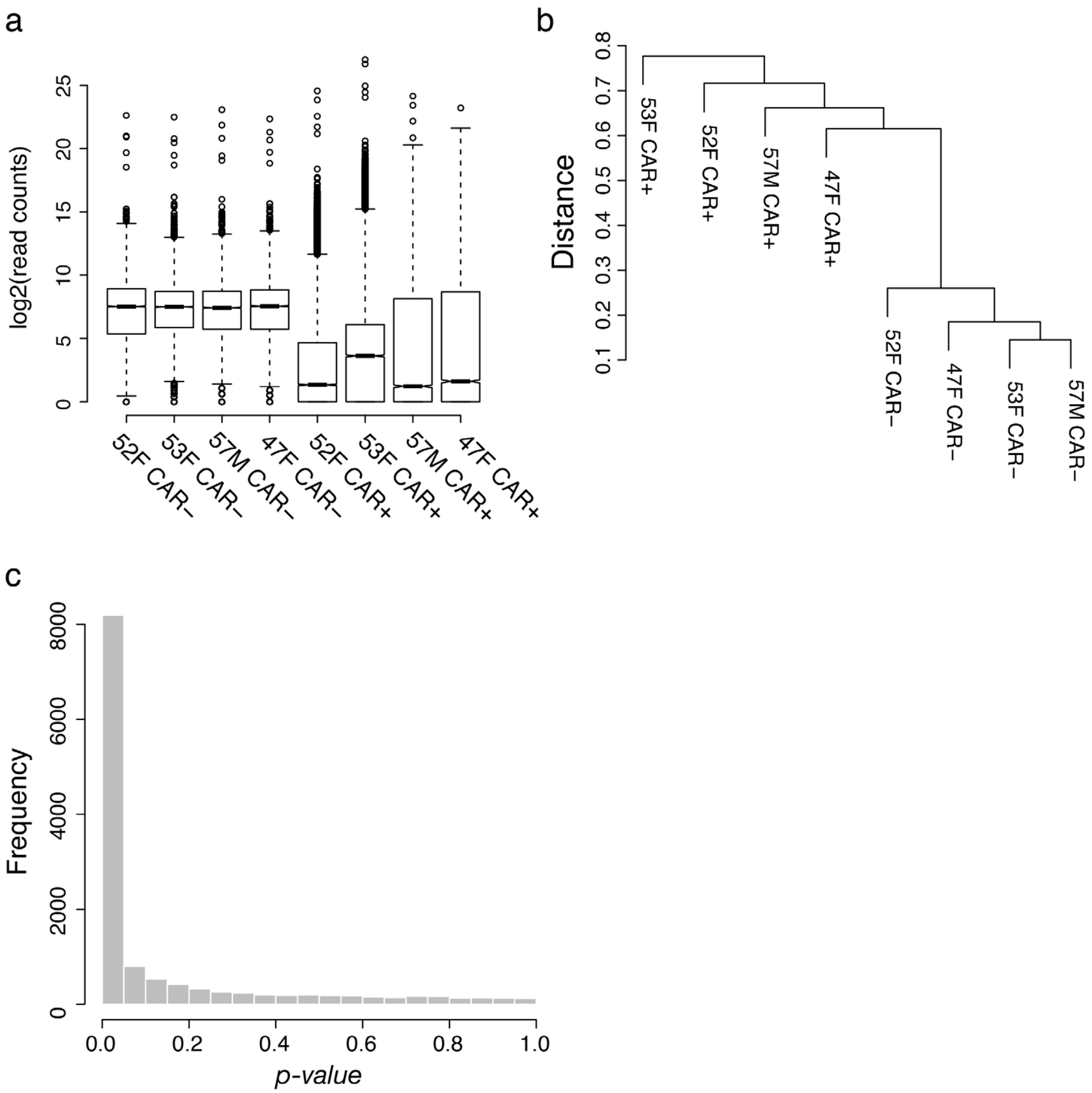


**Figure 3 - figure supplement 2: Quality control for adult human retinal FIN-Seq data.**

a) $Log_2$-transformed read distribution plot for sequenced human retinal samples. b) Dendrogram of read counts shows clustering of CAR$^{+}$ samples CAR$^{-}$ samples. c) A plot of frequencies of *p*-values shows an even distribution of null *p*-values.

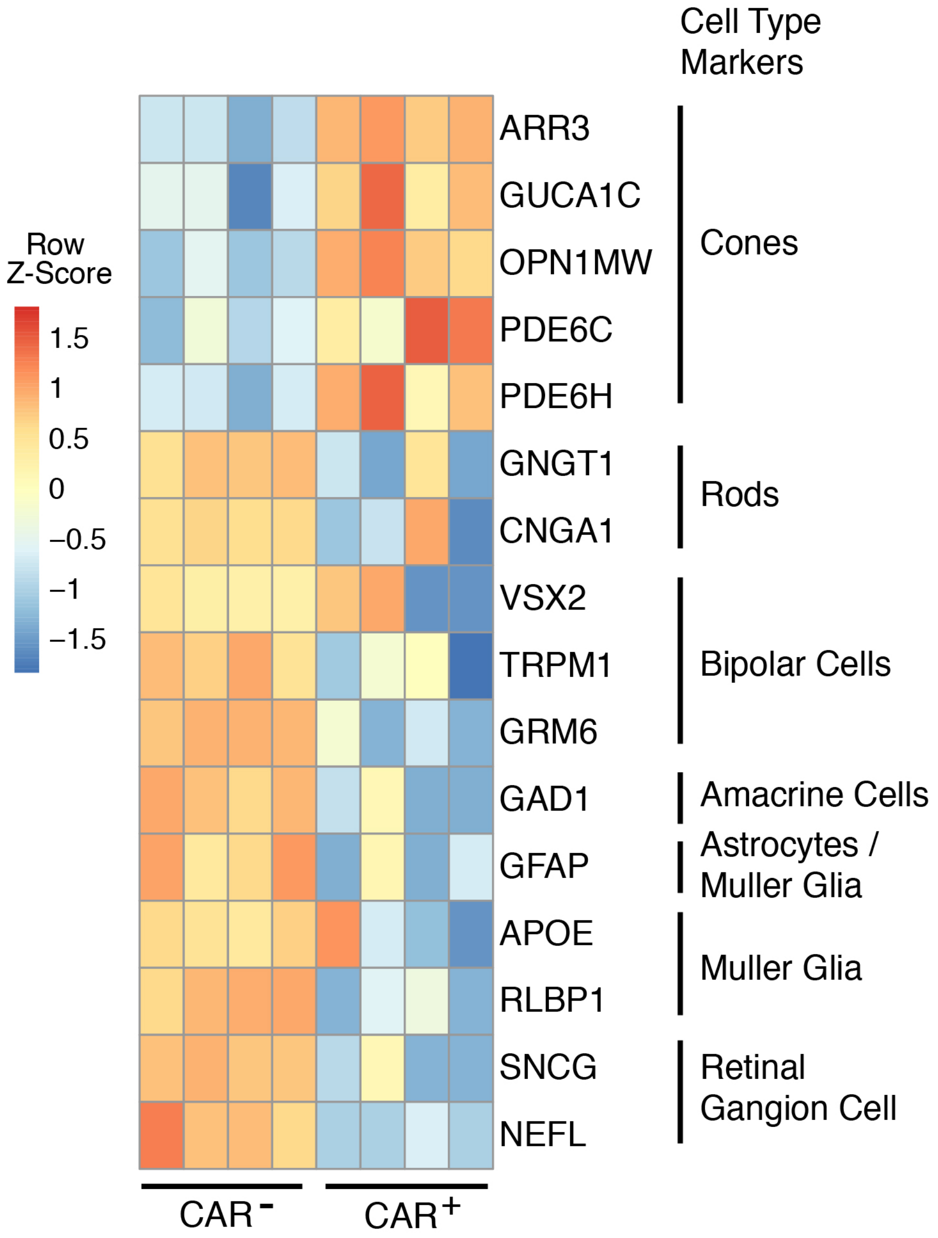


**Figure 3 - figure supplement 3: $CAR^+$ population contains mostly cone photoreceptors.**

A heatmap representing relative expression levels of human cell class specific markers previously identified by single cell RNA sequencing of a human retina sample (Lukowski et al., 2018). Markers of cone photoreceptors, which expresses CAR, were enriched in the $CAR^+$ population, indicating that this population contains mostly cone photoreceptors.
